# Ultrafast, alignment-free detection of repeat expansions in next-generation DNA and RNA sequencing data

**DOI:** 10.1101/2021.04.05.438449

**Authors:** L.G. Fearnley, M.F. Bennett, M. Bahlo

## Abstract

Short tandem repeat expansions are an established cause of diseases such as Huntington’s disease. Bioinformatic methods for detecting repeat expansions in short-read sequencing have revealed new repeat expansions in humans. Current bioinformatic methods to detect repeat expansions require alignment information to identify repetitive motif enrichment at genomic locations. We present superSTR, an ultrafast method that does not require alignment. We demonstrate superSTR’s ability to efficiently process both whole-genome and whole-exome sequencing data. Using superSTR we perform the first analysis of the UK Biobank to efficiently screen the exomes of 49,953 biobank participants for repeat expansions. We identify known mutations, as well as diseases not previously associated with REs. We further demonstrate the first bioinformatic screening of RNA sequencing data to detect repeat expansions in patients with spinocerebellar ataxia and Fuchs’ endothelial corneal dystrophy, and mouse models of myotonic dystrophy. superSTR is a highly computationally-efficient repeat expansion tool screening and detection tool for genomewide novel repeat expansion analysis, significantly outperforming existing methods. superSTR is available from https://github.com/bahlolab/superSTR.

## Introduction

Short tandem repeats (STRs) are nucleotide sequences comprised of repetitive motifs 1-6 nucleotides (nt) in length found within the genomes of all living organisms. They number in the tens of thousands and make up nearly two-thirds of a healthy human genome^1^, playing important roles in an array of biological functions^1–3^. Individual tandem repeats may vary in length and be subject to distinct selective pressures^4,5^. Variation in STR length has been attributed to events including strand-slippage during replication^6^, retrotransposition^7^, unequal crossing over in meiosis^8^, and DNA repair^9^.

Expansions of STRs are associated with a number of phenomena. Repeat expansions (REs) in genes which show increased variation in length between humans and other hominids are enriched for sensory and cognitive processes^10^. Some REs lead to the formation of fragile sites in the chromosome, resulting in increased rates of translocation and breakage^3^. This fragility may be beneficial, facilitating parallel adaptive evolution of certain traits^11^. REs are associated with a wide range of human diseases. Somatic REs are a hallmark of hereditary non-polyposis colorectal cancers^12^. Germline REs play causal roles in over fifty neurogenetic and movement disorders^4,13–15^. The presence of some germline expansions associated with neurodegenerative disorders also appears to reduce overall risk of other diseases, including certain cancers^16^. Taken as a whole, these phenomena suggest complex, potentially evolutionarily-conserved mechanisms of action.

Many biological mechanisms have been proposed as to how REs might impact cells beyond simple disruption of critical genomic sequence (Figure 1). REs may contain targets for DNA binding proteins^17^, or impact genomic regulatory elements^2^. REs can cause alteration of chromatin structure and conformation^18^, and impact methylation and other epigenetic mechanisms^19^. They may be transcribed, impacting RNA splicing by generating binding sites for RNA-binding proteins^20^, or affecting RNA degradation^21^. Transcribed REs may be translated, leading to protein aggregation^22^, accumulation of toxic polypeptides^23^, and disruption of cellular transport systems^24^.

**Figure 1:**
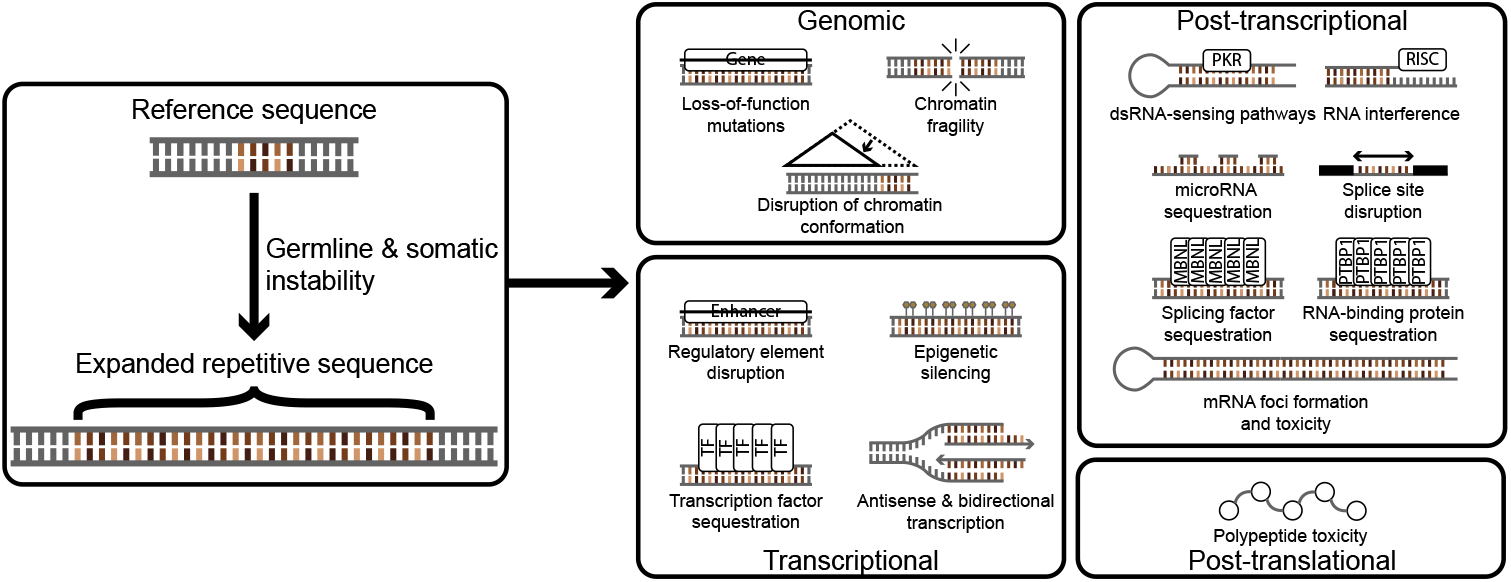
Consequences of short tandem repeat expansions. Presence of a short tandem repeat expansion may have structural or chemical consequences in addition to functional impact on genes, on the transcriptional capacity of the cell, in alteration of post-transcriptional and post-translational phenotypes, generation of cytotoxic products, or triggering of apoptosis.

*In vitro* assays by PCR and Southern blotting that target a specific motif or genomic locus are primarily used to investigate STRs in clinical contexts^25^. Short-read next-generation sequencing (NGS) based detection of REs is more common in a research context, and includes both whole-exome and whole-genome sequencing (WES and WGS). The first generation of bioinformatic RE detection methods for NGS data test known motifs at predefined genomic loci (STR loci) within an alignment^26–31^. These estimate the size of a putative repeat at each potential STR locus, and may proceed to test for statistical significance by case-control or outlier analysis^28,29^. Enumeration of STR loci in a reference is a separate bioinformatic challenge and uses tools such as Tandem Repeats Finder^32^.

Recent *de novo* methods perform RE discovery within alignments without prior knowledge of STR loci. These methods search for patterns of poor alignment to the reference genome, which are characteristic of REs. A *de novo* strategy used in discovery of the causal RE in cerebellar ataxia with neuropathy and bilateral vestibular areflexia syndrome (CANVAS)^33^ involved identifying candidate genomic regions via linkage analysis^34^, then visual inspection of aligned reads in these areas. Manual approaches are limited by the labour-intensiveness of the process and the number of samples needed for linkage analysis. ExpansionHunter Denovo (EHDN) automates *de novo* STR discovery by implementing heuristics on the mapping quality of alignments of reads to a reference^35^. EHDN was also used in the detection of CANVAS-associated REs^36^. A key limitation of all such methods lies in the alignment step; the choice of aligner and alignment parameters is critical and must be consistent for valid comparison across cohorts.

An alternative approach is to simply test all reads for repetition. This is implemented in TRhist and mreps^37,38^. Both methods use efficient algorithms based on string decompositions into unique, minimal representations, followed by steps that find maximal periodicities in the decomposed string^39,40^. Both methods are conceptually similar to tools used in assembled genomes^32^ but process many millions of short genomic sequences rather than comparatively small numbers of long genomic scaffolds. TRhist was used in the detection of REs associated with familial adult myoclonic epilepsy^41^, and neuronal intranuclear inclusion disease and oculopharyngodistal myopathy^42^. Sequential processing of all reads causes extremely long processing times when analysing data from NGS experiments with moderate target depth.

We present superSTR, the first method demonstrated to rapidly identify REs in population-scale cohorts without alignment or time-consuming processing of all reads in NGS experiments. superSTR uses a fast, compression-based estimator of the information complexity of individual reads to select and process only those reads likely to harbour expansions. We demonstrate superSTR’s ability to identify samples with REs and to screen motifs for expansion in raw sequencing data from short-read WGS experiments, in biobank-scale analysis, and for the first time in direct interrogation of repeat sequences in RNA-seq.

## Results

### Overview of superSTR – a method for identifying short repetitive elements in short-read data

superSTR efficiently identifies reads containing repetitive genomic sequence for processing by estimating the information content of each read using compression^43,44^. It relies on repetitive sequence being more compressible by zlib^43^ than non-repetitive sequence (Figure 2a). A ratio, *C*, is computed from the ratio of the size of the read in bytes after compression to its uncompressed size. This computation is extremely fast due to the highly optimised nature of the zlib library. We impose a threshold on C that triages up to 90% of WGS reads as non-informative.

**Figure 2:**
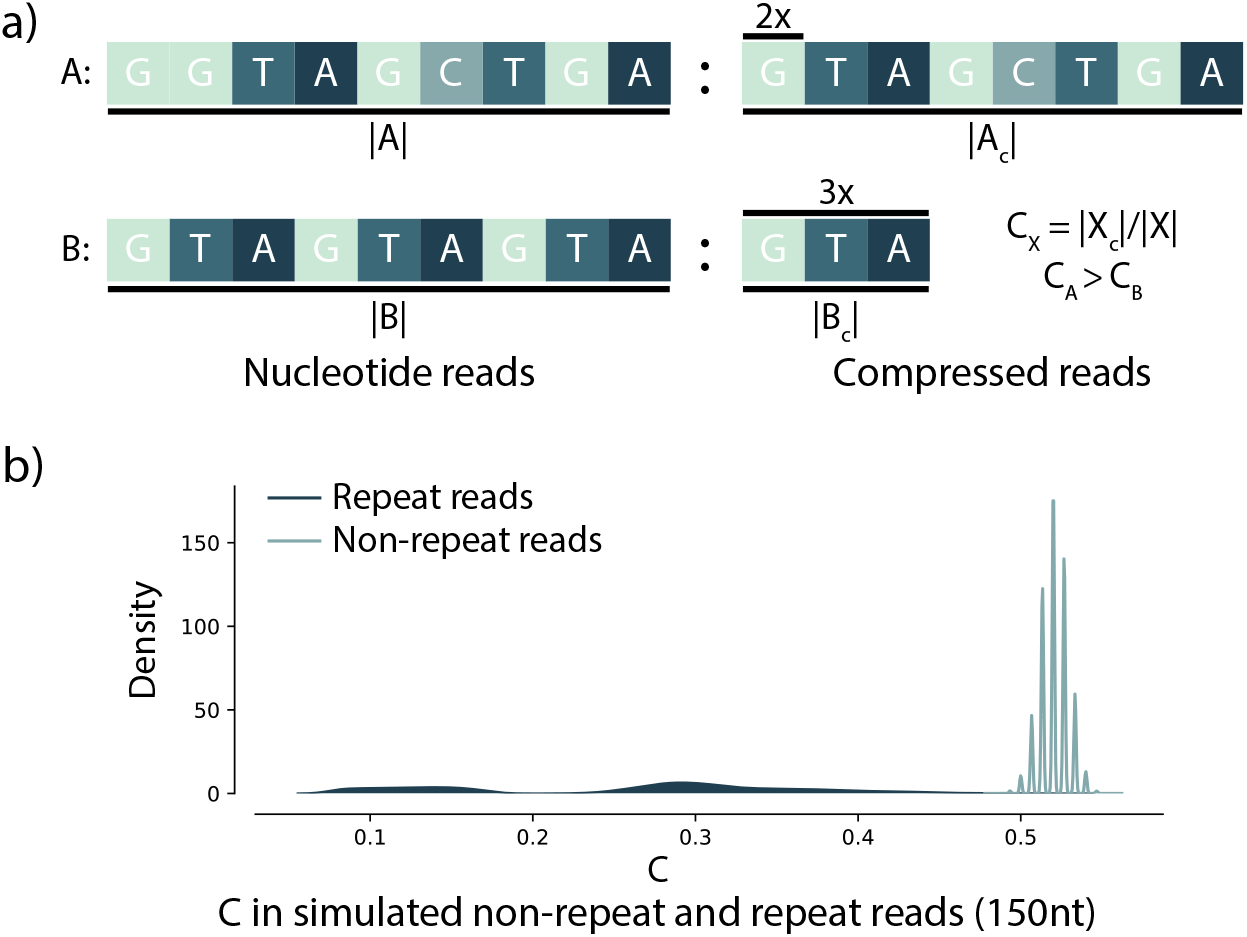
superSTR’s compression heuristic and its performance in simulated reads. a) superSTR relies on relative compressibility to distinguish between repeat containing reads. Compression with zlib involves removal of duplication. Repeat A (which does not contain significant repetition) will compress less than repeat B (which does), and the ratio of compressed size to uncompressed size will be greater for A than B. B) Distribution of C compression ratios in 150nt pseudorandom reads and repeat-containing reads drawn from a distribution where nucleotides are equiprobable and no errors are present. A more complete characterization across different distributions, read lengths and error rates is contained in Supplementary Figures S1-S5.

We evaluated *C* in classifying simulated repeat-containing reads against pseudorandom sequence drawn from four background nucleotide distributions. These were the uniform distribution, where each nucleotide is equiprobable, the genomic distributions of *Homo sapiens*, the GC-rich bacterium *Streptomyces coelicolor*, and the AT-rich intergenic regions of the protozoan *Plasmodium falciparum*. We tested *C* on common read lengths for experiments deposited in the NCBI Sequence Read Archive, noting that shorter read lengths are generally associated with older data sets. We randomly selected repeat motifs up to 15nt with all possible repeat lengths up to the read length (Supplementary Information).

A classifier using *C* to identify repeat-containing reads performs near-perfectly in simulations at read lengths greater than or equal to 75 nucleotides in length. *C* performance degrades, fewer reads are identified as non-informative, and the classifier becomes less specific as read lengths decrease below 75nt (Supplementary Information, Supplementary Figure S2). We further tested performance of *C* under varied sequencing error rates and in distinguishing simulated repeats from random sequence, observing similarly high performance (Figure 2b, Supplementary Figures S3—S5). We provide a set of optimal threshold values for *C* for the distributions and read lengths from these simulations (Supplementary Table 2).

Reads that pass the *C* classification step are processed using the maximal approximate periodicity algorithm as implemented in mreps^38^ to identify repetitive substrings within each read. We generate counts of the repeat length of each motif in each sample and calculate a combined information score which is then visualised and interrogated using statistical methods to detect REs (Methods).

superSTR is sensitive to the presence of background repetition in the genome, which can render outlier detection difficult if an expansion contains common repeating sequences such as telomeric repeat motifs. It is also unable to distinguish between complementary repeats if 5’-3’ orientation of the read is unknown. This is provided by most sequencing-by-synthesis methods.

### Detecting repetitive element outliers in short-read sequencing of individuals with known repeat expansions

We assayed 150nt paired-read whole-genome sequencing data from the Illumina Polaris dataset of 270 individuals. The Diversity cohort is 150 samples of unascertained RE status, and the Repeat Expansion (RE) cohort is 120 samples with orthogonally validated RE lengths^31^.

We performed motif screening across the Illumina Polaris WGS cohort (Figure 3a-e)). superSTR’s motif screening uses a one-sided Mann-Whitney U test to evaluate whether the distribution of information scores is the same in the test group and the Diversity cohort, or whether the test group distribution is greater. Statistical significance was estimated using permutation testing to generate *p*-values that were subsequently corrected for multiple testing using the Benjamini-Hochberg procedure^45^ (Methods). Permutation testing was used instead of the standard distribution for the Mann-Whitney U test due to some of the unequal variances observed.

**Figure 3:**
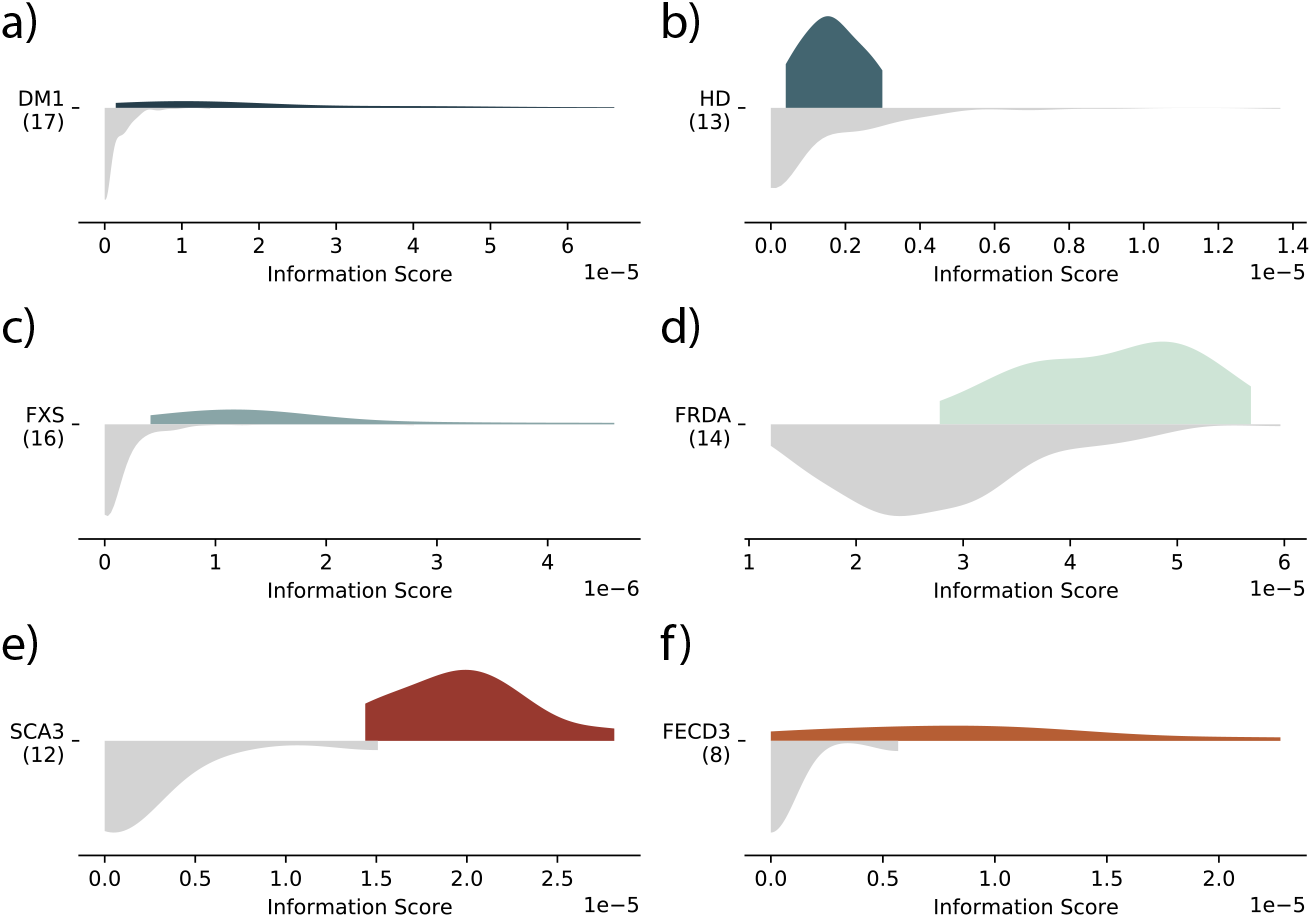
superSTR analysis of WGS and RNA-seq RE data. Controls are shown in grey (lower part of each graphic), and affected individuals in color (upper part of each graphic). a), b), c), and d) show comparison of groups within the Illumina RE cohort to the Illumina Diversity cohort. e) and f) show RNA-seq analysis. a) AGC profile of DM1-bearing individuals with long-tailed distribution characteristic of relatively large RE; b) AGC profile of HD-bearing individuals with a much shorter RE; c) CCG profile of FXS individuals; d) AAG profile of FRDA individuals. e) RNA-seq AGC profile of peripheral blood mononuclear cells from 12 individuals with SCA3 RE against 12 matched non-SCA3 controls. f) RNA-seq AGC profile of from eight patients with confirmed FECD3 expansions and six controls without FECD (of any type).

We detected 27 unique motifs with corrected *p*-values (*p*) < 0.05 in at least one Polaris RE group (Supplementary Data). We observed significant differences in AGC motif median difference in myotonic dystrophy, type 1 (DM1) (*p*=4.1 × 10^-3^) and in Huntington’s disease, which is associated with a smaller CAG expansion (*p*=3.8 × 10^-2^). Shifts in the AAG distribution were significant across the group of individuals with Friedreich’s ataxia (FRDA) and those with premutation-length FRDA alleles (*p*=4.1 × 10^-3^ and *p*=1.4 × 10^-2^ respectively). We observed significant shifts in the CCG information score in individuals with Fragile × Syndrome (FXS) and again in individuals with premutation-length FXS alleles (*p*=4.1 × 10^-3^ for both groups). Pathogenic AGC, AAG and CCG motifs were only significantly expanded in their associated disease groups. Among the 28 remaining motifs we note that the hexamer AGAGGG significantly differed in all seven groups; the next most common motif was AGGG, present in five (DM1, FRDA, HD, FXS Premutation and FXS Normal). Many of the repeats detected (including AGAGGG, AAGG, AGG, and AAGGG) have been associated with regions predicted to form G-quadruplex structures^46^, which play complex roles in neurodegenerative disorders^47^.

We identified individual samples with REs using a novelty detection approach reporting samples exceeding the BCa-bootstrapped estimate of the 95% confidence interval of the 95^th^ percentile of the information score in the Diversity cohort (Methods). Fourteen of 17 DM1 individuals in the Polaris cohort were called as outliers for the AGC motif using this method. Seven of 14 FRDA cases were called as AAG motif outliers, along with one carrier and a single individual who is likely a FRDA expansion carrier. 13 of 16 FXS cases were called as outliers for the CCG motif, along with 4 FXS premutation individuals, a non-affected relative of a FXS case, and a complex FXS case (NA07063).

Ranking samples by AGC information score revealed several individuals within the Diversity cohort with scores intermediate between DM1 individuals and individuals with smaller expansion disorders. EHDN analysis of these detected CAG expansions in *TCF4* (associated with Fuchs’ endothelial corneal dystrophy type 3 (FECD3)), an intronic CAG expansion in the *CA10* gene, and an apparent heterozygous expansion for spinocerebellar ataxia type 1 (SCA1) in *ATXN1* exceeding that disease’s pathogenic threshold. The finding of a SCA1 expansion was not supported by targeted-locus analysis using ExpansionHunter^31^, which assayed the locus as a homozygous expansion 6nt below the pathogenicity threshold.

### Motif screening in the UK Biobank

We obtained and analysed 75nt paired-read WES data for 49,953 individuals with GP and hospital-level clinical records mapped to ICD10 coding from the UK Biobank’s FE WES dataset^48^ as described in the Methods. We analysed differences in the information scores across tri-, tetra- and pentamer motifs within four-character UK Biobank ICD10 codes corresponding to diseases of the nervous system (Chapter VI, G00-G99) and eye and adnexa (Chapter VII, H00-H59) between individuals with that diagnosis and the rest of the Biobank. P-values were corrected within each ICD code using the Benjamini-Hochberg correction across motifs with non-zero information scores tested within each code (up to 149 motifs). Motifs were reported against an ICD code if they met a significance threshold of 0.05.

We identified 46 unique motifs comprising six trimers, 10 tetramers, and 30 pentamers in 60 disorders meeting these criteria (Table 1, Supplementary Data). The AGC motif information score distribution was significantly greater in individuals with myotonic disorders (G711, *n*=13, *p*=3.4×10^-3^) than those without, consistent with presence of DM1. Significant differences in AGC scores were also observed for individuals with hereditary corneal dystrophies (*n*=93, *p*=6.3 × 10^-3^), consistent with presence of FECD3. Follow-up analysis with EHDN localized these putative expansions to the expected *DMPK* and *TCF4* genes for 77% and 67% of hits respectively, along with a number of other loci (44 additional genes for G711, and 131 for H185). Among the genes to which EHDN mapped CAG repeats in individuals diagnosed with hereditary corneal dystrophies (H185) were a number of genes with non-RE mutations linked to ocular disorders and dystrophies. These genes and linked disorders included *AGBL1* (*n*=10, causal for FECD type 8^49^), *PDK3* (*n*=23, Charcot-Marie-Tooth disease, type 6, which involves development of optic atrophy among other symptoms^50^), *POLG* (*n*=2, progressive external ophthalmoplegia^51^), and *RAB28* (*n*=1, previously associated with cone-rod dystrophies^52^).

**Table 1:** superSTR analysis of UK Biobank data – significant trimers in motif screening of ICD10 codes and post-screening localization of superSTR leads by EHDN. Repeat sizes are as reported by EHDN and are listed for qualitative purposes only due to high uncertainty resulting from low coverage and numbers of supporting reads in some areas of the WES data. Highlighted genes have associations with related disorders. An extended version of the above table with complete gene lists incorporating data from tetramers and pentamers is included in Supplementary Data.

| ICD | ICD Description | # samples in cohort | Motif | MW statistic | FDR-corrected p-value | # samples with EHDN paired IRRs | EHDN localised genes (n, mean estimated het allele lengths) |
| --- | --- | --- | --- | --- | --- | --- | --- |
| H185 | H18.5 Hereditary corneal dystrophies | 93 | AGC | 3113774 | 0.0064 | 88 | **<u>TCF4 (63, 351.0), AGBL1 (10, 78.4), PDK3 (23, 99.95652173913044), POLG (2, 64.0), RAB28 (1, 74.0)</u>** & 127 other genes |
| G911 | G91.1 Obstructive hydrocephalus | 14 | AAC | 430504 | 0.035 | 1 | CHID1 (1, 97.0), TNFRSF19 (1, 62.0) |
| G711 | G71.1 Myotonic disorders | 13 | AGC | 530258 | 0.0034 | 13 | **<u>DMPK (10, 1736.7)</u>**, 44 additional genes (including MLTT3, TCF4, CA10, TBP) |
| G568 | G56.8 Other mononeuropathies of upper limb | 15 | AGG | 533234.5 | 0.038 | 0 | **<u>DCAF8L2 (7, 86.4), FGFR4 (2, 68.5), CHD3 (1, 61.0), EPHB3 (1, 61.0), GDNF (1, 45.0)</u>**, WIZ (1, 117.0), NAA38 (1, 61.0), _ERICH6 (1, 48.0) |
| G439 | G43.9 Migraine, unspecified | 2352 | ACG | 56157179 | 0.036 | 5 | **<u>CASZ1 (3, 62.0)</u>** |
| H654 | H65.4 Other chronic nonsuppurative otitis media | 95 | ACG | 2414382 | 0.023 | 1 | No localisation |
| G440 | G44.0 Cluster headache syndrome | 2082 | CCG | 51810209 | 0.045 | 113 | **<u>FMR1 (1044, 94)</u>**, AR (532,74) and 113 additional genes |
| G440 | G44.0 Cluster headache syndrome | 2082 | ACG | 50023184 | 0.020 | 5 | **<u>CASZ1 (3, 62.0)</u>** |
| H269 | H26.9 Cataract, unspecified | 2680 | ATC | 64490674 | 0.017 | 27 | CENPP (10, 58.8), ASPN (10, 58.8) and 41 other genes not previously linked to cataract disorders |
| G048 | G04.8 Other encephalitis, myelitis and encephalomyelitis | 17 | ACG | 448775 | 0.038 | 1 | No localisation |
| H210 | H21.0 Hyphaema | 29 | ATC | 849421 | 0.033 | 1 | ABCA8 (1, 168) |

We observed a statistically significant difference in the AGG motif for mononeuropathies of the upper limb (G568, *n*=15, *p*=3.8 × 10^-2^). Follow-up analysis of those individuals with EHDN detected anchored repeat reads in 13 individuals in *DCAF8L2* in 7 participants, *FGFR2* in 2 participants, and *EPHB3*, *ERICH6*, *CHD3*, *GDNF* and *WIZ* in 1 participant each. Five of these genes have links to neuropathies. *DCAF8* (a paralog of *DCAF8L2*) has been associated with giant axonal neuropathy 2^53^, *FGFR2* disruption leads to axonal neuropathy^54^, and *GDNF* reduces symptoms of neuropathy in mouse models^55^. The mouse ortholog of *CHD3* participates in the nucleosome remodeling and deacetylase chromatin remodeling complex (NuRD), a complex required for peripheral nerve myelination which causes neuropathy in mouse models when disrupted^56^. *EphB3*, another murine ortholog of a screen result, is involved in adult axonal plasticity and recovery from injury^57^.

We detected statistically significant differences in CCG motif information score for cluster headache syndrome sufferers (G440, *n*=2082, *p*=4.5 × 10^-2^). EHDN localized those reads to the Fragile X-associated *FMR1* locus for 50% of these individuals, with a mean expansion length of 94 repeats. FXS premutation length expansions have previously been associated with migraine headache in family studies of fragile-X associated tremor ataxia syndrome and in relatives of individuals diagnosed with FXS^58^.

The ACG motif is an exceptionally uncommon microsatellite in the human genome, appearing in 16 loci across the genome, only three of which appear in exons^59^. It also appears rarely in assayed UK Biobank WES samples. We were able to detect several highly significant differences in ACG distributions. A group of 11 individuals were diagnosed as having both migraine (G439, *n*=2352, *p*=3.6 × 10^-2^) and cluster headache syndrome (G440, *n*=2082, *p*=1.9 × 10^-2^). Analysis with EHDN localized repeats in the exomes of two of these individuals to the vicinity of exon 21 of *CASZ1. CASZ1* is a zinc finger transcription factor expressed during brain development^60^ in which a single nucleotide polymorphism has previously been associated with migraine by meta-analysis and GWAS with odds ratio [1.06-1.17]^61^.

### Direct detection of repetitive elements in RNA-seq data

We obtained and analysed publicly available RNA-seq data for experiments profiling expression in spinocerebellar ataxia type 3 (SCA3) and FECD3 patients, and from experiments using RNA-targeting CRISPR-Cas9 to clear CAG repeat-bearing RNA from cultured cells and mouse models.

#### Spinocerebellar ataxia type 3

SCA3 is caused by heterozygous expansion of CAG in *ATXN3*. Incomplete penetrance is observed above 44 repeats, and pathogenicity occurs when >52 repeats are present^62^. We analysed RNA-seq data of peripheral blood mononuclear cells (PBMCs) from 12 individuals with SCA3 and 12 age- and gender-matched control samples^63^. The AGC motif information score distribution was significantly greater than the control group (*p*=2.9×10^-3^, Figure 3e, Supplementary Data). Outlier detection identified all ten SCA3 cases as outlier samples against the control group.

#### Fuchs endothelial corneal dystrophy, type 3

FECD3 is a subtype of Fuchs endothelial corneal dystrophy (FECD) associated with an intronic heterozygous CTG expansion in *TCF4*. Presence of more than 40 repeats is associated with significantly increased risk of FECD^64^ but the expansion is incompletely penetrant. We obtained an RNA-seq dataset describing corneal endothelium of 12 eyes from 12 persons diagnosed with FECD undergoing transplantation and six controls of donor eyes unsuitable for transplant^65^. Eight of the FECD samples were confirmed to have the *TCF4* expansion by STR-targeting and triplet-primed PCR, and the remaining four determined to have nonexpansion FECD. *TCF4* expression is high in the corneal endothelium, and there are conflicting reports as to whether the presence of the CTG expansion increases expression of the *TCF4* mRNA^66–68^. We identified a significant increase in information score for AGC motifs in individuals with CAG expansions in *TCF4* (*p*=1.5×10^-2^, Figure 3f, Supplementary Data). Outlier detection identified five of eight individuals with *TCF4* expansions as outliers compared to controls.

### Myotonic dystrophy 1 and long CTG repeat cell line experiments

Myotonic dystrophy, type 1 (DM1) is caused by a CTG expansion in the 3’ untranslated region (UTR) of *DMPK*. The pathogenic range is broad, starting at 50 repeats for mildly-affected individuals and can exceed 2,000 repeats in individuals with severe disease. DM1 expansions induce formation of nuclear RNA foci that can be visually detected by fluorescent in situ hybridization (RNA FISH)^69^.

We analysed data from two studies with RNA-seq of cells treated with CUG-degrading dCas9^70,71^. The first study was performed in myotubes derived from induced pluripotent stem cells from a DM1 patient and an unaffected individual^70^. superSTR analysis was unable to detect differences between treatment and non-treatment groups (*p*=0.31 for the expected AGC motif, Supplementary Data). Investigation revealed the total absence of long AGC sequences in the RNAseq data despite the confirmation of their presence by RNA FISH. The second study was conducted in skeletal muscle from DM1 model HSA^LR^ mice that express >200 AGC repeats^71^. superSTR identified AGC as the sole outlier motif (*p*=1.2×10^-3^, Supplementary Data) in HSA^LR^ mice treated with non-targeting dCas9 (*n*=4) compared to the wild-type mice and HSA^LR^ mice treated with UGC-degrading Cas9 (*n*=16). Outlier detection based on information score correctly identified all four repeat-bearing HSA^LR^ mice as outliers with no false positives.

### Execution time and computational benchmarking

RE detection methods rely on statistical analysis of their output. These analyses are performed in two ways: (i) case control or motif screening analyses, where test statistics from individuals with possible REs are compared to a set of controls known to be free of REs, or (ii) outlier detection, where a set of samples of unascertained status likely containing a mix of REs and non-expanded sequence undergo analysis to detect outliers. The latter study design assumes a heterogeneous cohort where only a small proportion of the cohort harbour a particular RE, forming a potential group of outliers. If the outlier group is too large many of the statistical approaches will fail ^13,29^.

No statistical analysis methods are provided for TRhist; and as a result we were only able to evaluate its execution time, not its RE detection performance. We benchmarked TRhist, EHDN and superSTR using RE data from the Polaris dataset (150nt paired-end WGS data generated at a target sequencing depth of 30x coverage aligned to a GRCh38 reference) as described in the Supplementary Materials.

In testing on WGS data, superSTR processed an average of 444 million paired reads per individual within 4.5 hours (mean 258 minutes) using one CPU and less than 200MB of RAM.

TRhist execution times exceeded a cut-off at 36 hours. EHDN completed analysis with mean execution time of 24h 53m and maximum time of 30h 46m. EHDN execution times were dominated by the alignment step, which required up to 15 CPUs and 30GB of RAM at peak demand; EHDN consistently processed prepared alignments within an hour using a single CPU with peak memory demand of 2.5GB of RAM.

Both superSTR motif screening and EHDN case-control motif analysis identified statistically significant enrichment of the pathogenic ACG motifs in DM1, AAG motifs in FRDA, and CCG motifs in FXS. superSTR, but not EHDN, identified statistically significant enrichment of pathogenic AGC (CAG) motifs across the Huntington’s disease (HD) cohort. At the individual sample level, superSTR outlier detection consistently outperformed EHDN motif-level outlier analysis in identifying pathogenic samples. EHDN showed a recall of 0.53 for DM1-affected individuals called as AGC motif outliers compared to superSTR’s recall of 1.0; 0.75 for FXS-affected individuals (outliers for CCG) compared to 0.81, and 0.29 for FRDA-affected individuals (AAG outliers) compared to 0.50. A lack of orthogonal validation across all STR loci in the Polaris data prevents calculation of statistics requiring evidence of negative status, such as precision or F1-score. Neither superSTR nor EHDN identified HD individuals as AGC motif outliers. A detailed description of outlier calls at each sample is contained within the Supplementary Information.

superSTR and TRhist were computationally benchmarked on SCA3 RNA-seq samples described previously with an average library size of 15 million paired reads. EHDN does not support RNA-seq data. superSTR required mean execution time of 22m using 1 CPU and 5MB of RAM; TRhist required mean execution time of 4h 34m using 1 CPU and 2GB of RAM. Detail of benchmarking and results is provided in Supplementary Information and Supplementary Figures S6 and S7.

## Discussion

superSTR fundamentally differs from most existing bioinformatic STR detection methods in performing de novo searches for REs that requires neither alignment or specification of the locations of putative repeats. It produces motif-level predictions that are independent of alignment. superSTR furthermore sets up a statistical testing framework for motif screening and outlier detection, and provides an opportunity to detect REs in the many samples held in repositories such as dbGaP by enabling the rapid screening and re-screening of such data where computational resources are scarce. These data sources contain immense potential for discovery of both known, but undetected, as well as novel REs, an area of intense research interest.

The computational tool closest in capabilities to superSTR is the similarly alignment-free TRhist. superSTR substantially computationally outperforms that method by exploiting inherent properties of genomic reads, resulting in a 10-fold improvement in execution time. TRhist, a prototype RE analysis tool, has no statistical analysis framework to evaluate signals further preventing screening of biobank style cohorts.

superSTR outperforms the motif-based analysis of the alignment-dependent EHDN. This is likely due to a number of factors, one of which is the ability of superSTR to make use of signal from reads that only contain partial repeats. EHDN’s motif strategy analyses only pairs of reads that are mostly comprised of repeats with the same motif, and thus cannot be expected to detect expansions shorter than approximately the fragment length (twice the read length plus the insert size). As a result, superSTR can detect motif-level changes in shorter RE disorders such as Huntington’s disease. superSTR significantly outperforms EHDN for motif-based outlier detection due to fundamental differences in how the threshold for calling outliers is estimated. superSTR defaults to using the conservative upper 95% CI estimate of the 95^th^ quantile of the information score by the BCa bootstrap in only control samples with unascertained status. This behaviour is modifiable by the end-user through parameters. EHDN performs case resampling using both cases and controls using an assumption of a 5% outlier fraction. This assumption is violated in the Polaris cohort, and may also be violated in other cohorts for particular REs where it routinely generates high rates of false negatives at loci where repeats are relatively common, such as the FECD3-linked *TCF4* locus.

We recommend use of superSTR in a complementary fashion to existing RE detection tools as illustrated here with EHDN. superSTR performs best where the repeat expansion is longer than the read length, and on motifs not common in the genome. Alignment-based methods currently outperform superSTR when this is not the case, but their performance gains are balanced by the substantially increased computational cost associated with alignment and the significant multiple testing burden created in testing multiple loci per motif across large sets of diseases in cohorts such as the UK Biobank.

Our results in the UK Biobank motif screening analysis recover known REs and identify possible novel REs in genes linked to disorders not previously known to involve REs. We caution that there are many significant sources of confounding within the UK Biobank, and that further work is needed to interpret the results presented here, especially since many participants have only partial clinical data. The use of ICD10 categories as labels remains challenging when investigating specific diseases bundled with others in a single overarching category. There is also a suggestion of structure or underdiagnosis within the UK Biobank – FECD3 prevalence is estimated as 3-11% in Caucasians^72^, yet the prevalence of the ICD code representing *all* hereditary corneal dystrophies in the analysed cohort is 0.19%. As such our findings in a general cohort require validation in more disease-specific cohorts.

superSTR can directly analyse repeats in RNA-seq data without alignment to reference transcriptomes or genomes. Prior studies have largely focused on detection of downstream transcriptional signatures of expansion rather than direct detection of expanded sequence itself. There are several potential complications to such analysis. REs contained in intronic and exonic regions will likely show tissue-specific patterns of expression similar to those of the genes in which they are contained. Intergenic REs may not be expressed at all, although recent work shows that transcription initiation at microsatellites is much more widespread than previously thought^73^. Choice of RNA-seq library preparation method also appears to impact the viability of STR analysis. We observe some evidence of this complexity in DM1-derived myotubes, where the repeat is absent despite confirmation of its presence by RNA FISH imaging. These behaviours may require consideration given the emergent use of cell lines in investigating RE disorders and potential treatments.

## Methods

### Genome sequence data

The Illumina Polaris cohort consists of 270 samples, being 150 publicly available wholegenome sequences of individuals with unknown RE status (European Nucleotide Archive accession PRJEB20654), and a set of 120 whole-genome sequences from the Coriell Biobank with orthogonally validated pathogenic REs of varying lengths across eight disorders associated with three motifs (European Genome-Phenome Archive accession EGAD00001003562). The latter data is available on application as detailed in the European Genome Archive. Samples for both cohorts were prepared with TruSeq PCR-free sample preparation, then sequenced by an Illumina HiSeq × with 150×150nt paired read lengths to a target whole genome coverage depth of 30x. This dataset has been previously used to evaluate performance of REs detection methods^26,29,30,35^.

Whole-exome sequences of 49,953 individuals with unascertained RE status and available linked primary care records were obtained from the UK Biobank WES FE dataset in CRAM format (Data Fields 23163, 23164). These individuals were sequenced with the DT xGen Exome Research Panel v1.0, targeting a total 39 million base pairs of the genome (19,396 genes). Samples were sequenced with 75×75nt paired-end reads on the Illumina NovaSeq 6000 platform to a target 20x coverage depth. Primary care data containing readv2 and readv3 codes were converted to ICD-10 codes using the UK Biobank-supplied tables (UK Biobank Data-Coding 19), and those codes mapping to nervous system disease (G00-99) and diseases of the eye and adnexa (H00-H59) were tested.

### Transcriptome sequence data

All transcriptomic data is sourced from published experiments that are publicly accessible within the NCBI Sequence Read Archive; we briefly summarise these here.

RNA-seq data for SCA3 was obtained from a study describing the transcriptome of peripheral blood mononuclear cells in 12 SCA3-affected individuals with 12 age- and gender-matched controls from a study of patients at Xiangya Hospital in Changsha, China^63^ (SRA accession SRP168964). Samples were depleted of ribosomal RNA, cDNA synthesised using the TruSeq Stranded Kit, then sequenced on an Illumina HiSeq × Ten instrument with 100nt long reads.

RNA-seq data for FECD3 was obtained from a study conducted at the S. Fyodorov Eye Microsurgery Federal State Institution, Russia^65^ (SRA accession SRP186882). This study describes the transcriptome of the corneal endothelium of twelve individuals with FECD undergoing corneal transplantation, along with six control samples obtained from deceased individuals. Eight of these samples showed expansion in the *TCF4* gene when assayed using a triplet-primed polymerase chain reaction. Samples were depleted of ribosomal RNA and sequenced with 2×125 cycles on an Illumina HiSeq 2500 instrument. Mean read lengths of the data deposited in the Sequence Read Archive for this experiment varied among samples from 132nt to 124nt. We trimmed reads in this experiment to a uniform 124nt length prior to processing with superSTR.

RNA-seq data for DM1 was obtained from two studies of clearance of CTG-repeat RNA foci by RNA-targeted CRISPR-dCas9. The first study was conducted in human myoblasts derived from induced pluripotent stem cells that were in turn derived from a patient with DM1 (SRA accession SRP111361)^70^. These cells have a CAG repeat length estimated by multiple orthogonal methods as being between 1,793-2,700 repeats in the *DMPK* gene^74^. This study used Illumina TruSeq PolyA library preparation, and samples were sequenced on an Illumina HiSeq 4000 instrument, with a uniform 99×99nt read length

RNA-seq data for DM1 in a mouse model was obtained from a study of clearance of CTG-repeat RNA foci in skeletal muscle of HSA^LR^ mouse, which expresses 250 CTG repeats associated with the human skeletal actin promoter, and has 20-25 CTG repeats in the mouse *Dmpk* gene itself (SRA accession SRP266474)^71^. This study used Illumina TruSeq PolyA library preparation, and samples were sequenced on an Illumina HiSeq 4000 instrument. The SRA-deposited RNA-seq data for this experiment also varied in average spot length, with common read lengths in wild-type mice of 150×150nt paired reads and 100×100nt paired reads in the DM1-affected mice. We cropped reads in this experiment to a uniform 100nt length prior to processing with superSTR. We assigned ‘expanded’ status to runs of samples from untreated HSA^LR^ mice in this experiment (n = 8 runs) and ‘contracted’ status to treated HSA^LR^ mice and all WT mice in the experiment (n = 12 runs).

### superSTR algorithm - read processing

The sequence processing component of superSTR is implemented as a single-threaded application in the C programming language. Post-processing, outlier detection, and visualisation scripts are implemented in Python 3. Sequencing data is either read through the htslib library (v. 1.9) or streamed directly from file, pipe or standard input (for example, when reading sequence output piped to the software directly from the SRA Toolkit^75^). Each read is compressed using zlib^43^, set for the fastest possible compression speed. *C* is computed as the ratio of the size in bytes of the output data to the size in bytes of the input data (Supplementary Information). Thresholds on *C* are supplied by the user from either the pre-provided threshold tables or user-run simulation; we used the precomputed values for each analysis here. Reads passing this threshold are then analysed using the Kolpakov-Kucherov algorithm^40^ as implemented in a modified version of the mreps software with optimisation and modifications that enable its use in this context^38^. Each read identified as containing repetitive elements has their start and end location, motif, length and purity recorded.

### superSTR algorithm – post-processing and outlier detection

Post-processing steps summarise the output of the read processing methods into a list of the number of repetitive elements of each length (up to the read length) with a specified motif for each sample. This number is normalised by library size, and per-sample and per-motif files are generated. Summarisation and subsequent visualisation are performed in Python 3.8 code. An overview of the method is outlined in pseudocode in the Supplementary Information.

Summarisation produces a repeat count vector *ν* for each motif *m*, *ν* = *ν*_1_, *ν*_2_… *ν_j_*, where *ν_n_* is the number of times a repetitive sequence of length *n* was encountered in the sequencing data. The information score is then computed for each motif as & = (∑_*i*≤*l*≤*j*_ *r_l_* × *l*)/*n*, where *n* is the number of reads in the sequencing experiment. The default setting used in this study for *j* is the read length and *i* is [*j* × 0.75]. *i* and *j* may also be manually set as parameters, which permit detection of some shorter REs. superSTR handles cohorts where samples have been sequenced using mixed read lengths by trimming reads to a consistent length, as described in the RNA-seq section of these methods.

### Motif-screening and outlier analysis

We performed motif screening analysis using a one-sided Mann-Whitney U test as implemented in the scipy library^76^. The test was performed using the alternative hypothesis that randomly selected samples of interest would have larger information scores than samples from the background. We computed the p-value of test statistics by permutation testing of each motif-sample pair (10,000 permutations). We computed the reported permutation p-value with approximation to the exact value^77^ using the arbitrary-precision mpmath library^78^ to perform Gauss-Legendre quadrature. P-values were then false discovery rate-corrected using the Benjamini-Hochberg procedure as implemented in the statsmodels library^79^. Motifs were reported as significant if the corrected Mann-Whitney permutation p-value was less than 0.05.

superSTR’s default general-purpose outlier detection procedure is a one-versus-many strategy. The 95% confidence intervals for the 95^th^ percentile of each motif’s information score are estimated in a background or control cohort using the BCa-bootstrap^80^ as implemented in the arch package (version 4.19)^81^, and values that exceed the upper 95% confidence interval of the estimate of this percentile are reported as outliers.

### Benchmarking

We benchmarked the computational performance of superSTR for WGS and RNA-seq on FASTQ input against TRhist and ExpansionHunter Denovo 0.9.0 (including the alignment, sorting and indexing steps performed with BWA v0.7.17 and samtools v1.11). Execution time was benchmarked on four WGS and four RNA-seq samples on the Walter and Eliza Hall Institute of Medical Research’s HPC system (Intel Xeon Sandy Bridge E5-2690 v4 CPUs with data stored on a Quantum Xcellis Storage Appliance). Accuracy was benchmarked using the Illumina Polaris dataset, testing disease groups within the Repeat Expansion cohort against the ‘control’ population of the Diversity cohort. Full details of benchmarking are contained in the Supplementary Information.

## Acknowledgements

L.G. Fearnley is supported by the DHB Foundation Centenary Postdoctoral Fellowship in Neurogenetic Systems Biology. M. Bahlo is funded by an NHMRC Senior Research Fellowship (ID: 1102971). M.F. Bennett is supported by a Taking Flight Award from CURE Epilepsy. This work is supported by the Victorian State Government Operational Infrastructure Support and the Australian Government National Health and Medical Research Council Independent Research Institute Infrastructure Support Scheme. This research has been conducted using the UK Biobank Resource under Project 36610, “Exploring the genetic basis of healthy development and ageing: gene discovery and methods development.”.

