## Supplementary Information for "Ultrafast, alignment-free detection of repeat expansions in next-generation DNA and RNA sequencing data"

### Supplementary methods

*superSTR algorithm and compression heuristic*

The superSTR algorithm is as follows:

**Algorithm 1** superSTR: read-processing algorithm

**procedure** PROCESS_READS($sequences$, $threshold$) ▷ Note 1
  **for** $sequence \in sequences$ **do**
    $sequence_c \leftarrow$ zlib-compressed $sequence$
    $C \leftarrow |sequence_c|/|sequence|$ ▷ Note 2
    **if** $C \leq threshold$ **then**
      Find all maximal sequence repeats ▷ Note 3
      Output repeats for this read
    **end if**
  **end for**
**end procedure**

**Note 1:** *threshold* determination is described in the following sections.
**Note 2:** $|sequence|$ and $|sequence_c|$ are the in-memory sizes of *sequence* and $sequence_c$ in bytes.
**Note 3:** These steps are implemented using the mreps algorithm and software described in Kolpakov, R., Bana, G., Kucherov, G. (2003).

superSTR's compression heuristic, *C*, relies on repeat-containing reads having a lower information-theoretic complexity than non-repetitive strings. Various estimators of information-theoretic complexity have previously been applied to analysis of DNA in a number of contexts including (non-exhaustively) in detection of coding regions[1], alignment-free sequence similarity[2], and in tandem repeat detection in assembled genomes[3], and in a number of problems involving analysis of sequence in biological contexts[4]. Information-theoretic complexity is a field with a deep theoretical underpinning; we treat this informally here and refer the reader to work by Li and Vitanyi[5] and to Lempel and Ziv[6] for more-thorough treatment of Kolmogorov complexity and Lempel-Ziv complexity, respectively.

The complexity metric used in superSTR is the compression ratio of each read when encoded by the zlib algorithm, which uses the DEFLATE compression method[7,8]. zlib achieves compression by removal of duplicate substrings and their replacement with pointers that require less space, and by replacement of symbols based on frequency of use. It is the former element of the compression algorithm that allows superSTR to differentiate between repetitive and non-repetitive reads. A string containing repetition is more compressible by the DEFLATE algorithm than a string that contains no repetition. This heuristic holds in the presence of impurities in the repetitive sequence or errors in the sequencing process. However, as read lengths decrease, the performance of the compression heuristic decreases (as shown in Supplementary Figure S2). The chances of encountering duplicate substrings within the read decrease with read length, and the string becomes less compressible.

Other lossless compression methods should work in a similar fashion provided that they exploit the space reduction afforded by removal of duplicate substrings.

It is worth noting that zlib compression may behave counterintuitively when provided small, non-repetitive inputs. In this case, the compressed string may be larger than the uncompressed string. This occurs because in the worst case zlib will store uncompressed data in (up to) 16KB blocks. It requires 6 bytes of space for a header, then 5 bytes of block header per block. A 32-character string representing a 32nt read will have a size of 32 bytes; its zlib-encoded size in the worst-case is around 43 bytes, yielding a compression heuristic value of 1.34. Repetitive reads are still somewhat separable from non-repetitive reads in the context of short read lengths (as discussed in the following section), but use of superSTR or compression-based heuristics in these contexts requires particular care.

*Performance of the superSTR heuristic in simulated reads*

We simulated reads to test the superSTR heuristic by drawing strings of fixed length from a background nucleotide distribution. We generated 300,000 random strings drawn from the background to represent reads describing non-repetitive DNA, and 300,000 repeat-containing strings, comprised of 100,000 pure repeat reads, 100,000 reads which contained a repetitive prefix or suffix, and 100,000 reads which contained a repetitive sequence in the middle of the read sequence. Each repeat-containing string was generated with a randomly selected motif between 1 nucleotide (nt) and 15nt in length.

We first tested a set of four distributions selected to test the performance of the heuristic in GC-rich contexts (*Streptomyces coelicolor*), AT-rich contexts (*Plasmodium falciparum*), a context approximating the human genome, and the worst-case context where nucleotides were equally probable (Supplementary Table 1). Supplementary Figure S1 shows the distribution of *C* and separability of pseudorandom and repeat-containing reads for each of the four different nucleotide distributions.

We then repeated this process while varying the read length to reflect the read lengths common in the NCBI SRA; 32, 75, 150, and 300 nucleotides. Shorter read lengths (below 36nt) are typically associated with older experiments or non-mammalian data; we dropped simulation of repetition in the middle of reads in read lengths < 75nt due to the limited number of repeats possible. The distributions of compression heuristic for repeat-containing reads was non-overlapping with the distribution of the heuristic in pseudorandom sequence across these variations in read length (Supplementary Figure S2). We observed failure to compress pseudorandom sequence at the 32nt read length as expected and for the reasons outlined in the preceding section.

We then fixed the read length and repeated the process of performance evaluation with impure repeat sequences. Error rates of 0.0, 0.025, 0.05, 0.075 and 0.10 were selected; compression heuristic values increased but separability of repeat-containing and non-repeat containing reads was maintained across this range (Supplementary Figure S3).

Receiver operating characteristic and precision-recall curves were generated for read lengths of 75nt and 150nt for the uniform nucleotide distribution, *H. sapiens*, *S. coelicolor*, and *P. falciparum* (Supplementary Figure S4). Areas under receiver-operating characteristic (ROC) and precision-recall curve (PRC) was greater than 0.99 for all evaluations. We then proceeded to conduct an extended evaluation with a view to identifying the point where the ROC and PRC degraded; we observed consistent degradation of the performance of the compression heuristic as classifier in the 16nt read length range.

*Determining thresholds for superSTR analysis*

We recommend the use of a simulation approach as outlined above to elucidate the optimal threshold for analysis with superSTR. We provide code (in the simulation.py script) which computes the ROC and sum of precision-recall (PR) of the compression across the range of threshold values in a distribution, then provides the threshold which produces the maximum ROC and PR values as a conservative default setting. A set of precomputed values for these thresholds is available at the superSTR git repository and in Supplementary Table 2.

A more-aggressive threshold yields improved runtimes at the cost of omitting repeat-containing reads with short runs of repetitive sequence.

*Benchmarking – WGS and RNA-seq*

Benchmarking was conducted using servers in a high-performance computing (HPC) facility at the Walter and Eliza Hall Institute (WEHI) in Melbourne, Australia. We used human-derived data available from the NCBI SRA without restriction, with the exception of the two human WGS RE samples. These data are available on application to the data access committee (details of which are available at the European Genome-Phenome Archive, accession EGAD00001003562).

All NCBI SRA data was downloaded and processed using sratools version 2.10.9. Data was prefetched from the NCBI SRA, converted into FASTQ files using the fasterq-dump tool, then compressed using gzip.

<u>WGS data:</u> We benchmarked the performance of superSTR against the alignment-dependent de novo STR profiler ExpansionHunter deNovo 0.9.0 (EHDN)[9] and TRhist 1.0.1 on whole genome sequencing data.

We used the 1000 Genomes GRCh38 reference genome, which contains alt contigs, decoy and EBV sequences from the Genome Reference Consortium build 38 patch 13 (GCA_000001405), along with HLA sequences generated by Heng Li. This reference is available at:

ftp://ftp.1000genomes.ebi.ac.uk/vol1/ftp/technical/reference/GRCh38_reference_genome
/GRCh38_full_analysis_set_plus_decoy_hla.fa

For computational performance testing, two samples with unascertained RE status were randomly selected from the Illumina Polaris sequencing dataset of individuals from the 1000 Genomes cohort. The selected samples were HG01625, an Iberian Spanish male, and HG00449, a Southern Han Chinese female. The corresponding SRA accessions for sequence data were ERR1955416 and ERR1955537, respectively. Both WGS samples were sequenced using a PCR-free library preparation and 151x151nt paired-end sequencing on an Illumina HiSeq X Ten instrument[10].

Two publicly available samples with known RE expansion were selected from WGS samples[11]. The selected samples were Sample 9, an individual positive for DM1, and Sample 3, an individual positive for SCA3. The corresponding SRA accessions for these samples were SRR7205168 and SRR7205176, respectively. Both samples were sequenced using PCR-free library preparation on an Illumina HiSeq X Ten instrument[12].

Comparison of software outputs was performed using samples testing samples from the controlled-access Illumina Polaris Repeat Expansion cohort against the open-access Illumina Polaris Diversity cohort as controls. These WGS samples were sequenced using a PCR-free library preparation and 151x151nt paired-end sequencing on an Illumina HiSeq X Ten instrument[10].

WGS - Benchmarking procedure: superSTR takes as input unaligned fastq files as input. EHDN (and other alignment-dependent methods) take aligned reads in BAM or CRAM formats as input; we report the time for execution from unaligned FASTQ (including the necessary alignment step) and also the time for execution when provided with a BAM file.

Benchmarking was performed using the built-in execution and timeline reporting methods of the Nextflow workflow engine (version 20.10.0). We benchmarked all software from input FASTQ files; a breakdown of steps within the EHDN pipeline describes time required for alignment, sorting, indexing and processing with EHDN

**TRhist:** TRhist was run using the following command:

```
java -jar TRhist.jar -p -z -output_fasta out.fa input_R1.fastq.gz
input_R2.fastq.gz
```

**superSTR:** superSTR was run using the following command:

```
superstr --mode=fastq -t 0.49 -o input_R1.fastq.gz
input_R2.fastq.gz
```

**ExpansionHunter Denovo:** Alignment was performed using bwa 0.7.17r1188 to align reads to the 1000 Genomes GRCh38 reference genome with 16 threads, the resulting SAM file was coordinate sorted using samtools, again using 16 threads and the resulting bam indexed. ExpansionHunter denovo (EHDN) was run using the default settings.

The benchmarking ran the following commands:

```
bwa mem -Y -K 100000000 -t 16 GRCh38.fa input_R1.fastq.gz
input_R2.fastq.gz > aligned.sam

samtools sort -@ 16 -o aligned.bam aligned.sam

samtools index aligned.bam

ExpansionHunterDenovo profile --reads aligned.bam --reference
GRCh38.fasta --min-anchor-mapq 50 --max-irr-mapq 40
```

Case-control analysis: Case-control analysis for samples from individuals with diagnosis of Huntington's Disease (HD), Friedreich's Ataxia (FRDA), myotonic dystrophy 1 (DM1), and Fragile X Syndrome (FXS) was performed using default parameters and arguments for EHDN. superSTR analysis was performed using defaults for the information score metric (calculated on repeat lengths from 112-150nt); as a result, the calculated information score somewhat exceeds the lower bound for the pathogenic threshold for Huntington's disease; a custom analysis targeting this range is possible but was not performed in this benchmarking. TRhist does not provide any implementation of a statistical analysis.

Outlier detection: Outlier detection was performed in both superSTR and in EHDN by using both tools' outlier testing methods. All samples contained within the repeat expansion cohort of 120 samples were presented to the software in a single batch. EHDN motif-based outlier detection was performed using the method's default parameters. superSTR detection was performed with default parameters for superSTR's information score metric (repeat lengths 112-150nt; upper 95% confidence interval estimated by the BCa[13] bootstrap of the 95$^{th}$ quantile of the score in controls only, representing a 5% outlier fraction in controls).

RNA-seq: A set of four samples were selected from the RNA-seq produced for a study of spinocerebellar ataxia type 3 in patients from mainland China[12]. We selected two samples with known SCA3 expansion (SRR8195657, SRR8195661) and two without known expansion (SRR8195668, SRR8195670). All samples were sequenced using an Illumina HiSeq X Ten with random PCR library selection and 100nt single read sequencing[12].

RNA-seq benchmarking procedure: ExpansionHunter Denovo does not support RNAseq. We benchmarked computational performance of superSTR and TRhist.

**TRhist:** TRhist was run using the following command:

```
java -jar TRhist.jar -p -z -output_fasta out.fa input_R1.fastq.gz
input_R2.fastq.gz
```

**superSTR:** superSTR was run using the following command:

```
superstr --mode=fastq -t 0.49 -o input_R1.fastq.gz
input_R2.fastq.gz
```

*Benchmarking results*

Benchmarking results for computational performance on WGS data are presented in Supplementary Figure S6. Outlier calls for each sample in the RE cohort are presented in Supplementary Table 2 and the recall statistic is reported in the main text. RNA-seq benchmarking results for computational performance are presented in Supplementary Figure S7. All benchmarking results were generated in run reports from the Nextflow execution pipeline. Comparison of software outputs was not possible, as no implementation of a statistical analysis for motif detection is available for TRhist.

*Supplementary Material References*

1. Orlov, Y. L. & Potapov, V. N. Complexity: An internet resource for analysis of DNA sequence complexity. *Nucleic Acids Res.* (2004) doi:10.1093/nar/gkh466.
2. Ferragina, P., Giancarlo, R., Greco, V., Manzini, G. & Valiente, G. Compression-based classification of biological sequences and structures via the Universal Similarity Metric: Experimental assessment. *BMC Bioinformatics* **8**, 252 (2007).
3. Gusev, V. D., Nemytikova, L. A. & Chuzhanova, N. A. On the complexity measures of genetic sequences. *Bioinformatics* **15**, 994–999 (1999).
4. Aboy, M., Hornero, R., Abásolo, D. & Álvarez, D. Interpretation of the Lempel-Ziv complexity measure in the context of biomedical signal analysis. *IEEE Trans. Biomed. Eng.* **53**, 2282–2288 (2006).
5. Li, M. & Vitányi, P. *An Introduction to Kolmogorov Complexity and Its Applications*. (Springer New York, 1993).
6. Lempel, A. & Ziv, J. On the Complexity of Finite Sequences. *IEEE Trans. Inf. Theory* (1976) doi:10.1109/TIT.1976.1055501.
7. Deutsch, P. & Gailly, J.-L. RFC 1950 - ZLIB Compressed Data Format. *IETF* **RFC 1950**, 1–10 (1996).
8. Deutsch, P. RFC 1951 - DEFLATE Compressed Data Format Specification version 1 . 3 IESG. *IETF* **RFC 1951**, 1–15 (1996).
9. Dolzhenko, E. *et al.* ExpansionHunter Denovo: A computational method for locating known and novel repeat expansions in short-read sequencing data. *Genome Biol.* **21**, 102 (2020).
10. Dolzhenko, E. *et al.* ExpansionHunter: A sequence-graph-based tool to analyze variation in short tandem repeat regions. *Bioinformatics* **35**, 4754–4756 (2019).
11. Dashnow, H. *et al.* STRetch: Detecting and discovering pathogenic short tandem repeat expansions. *Genome Biol.* **19**, (2018).
12. Li, T. *et al.* RNA Expression Profile and Potential Biomarkers in Patients With Spinocerebellar Ataxia Type 3 From Mainland China. *Front. Genet.* **10**, 566 (2019).
13. Efron, B. Better bootstrap confidence intervals. *J. Am. Stat. Assoc.* **82**, 171–185 (1987).
14. Piovesan, A. *et al.* On the length, weight and GC content of the human genome. *BMC Res. Notes* **12**, 106 (2019).
15. Bentley, S. D. *et al.* Complete genome sequence of the model actinomycete Streptomyces coelicolor A3(2). *Nature* **417**, 141–147 (2002).
16. Hamilton, W. L. *et al.* Extreme mutation bias and high AT content in Plasmodium falciparum. *Nucleic Acids Res.* **45**, 1889–1901 (2017).

| | Nucleotide fraction | | | |
|---|---|---|---|---|
| | **G** | **C** | **A** | **T** |
| *Homo sapiens* [14] | 0.20435 | 0.20435 | 0.29565 | 0.29565 |
| *Streptomyces coelicolor* [15] | 0.3606 | 0.3606 | 0.1394 | 0.1394 |
| *Plasmodium falciparum* [16] | 0.097 | 0.097 | 0.403 | 0.403 |
| Uniform distribution | 0.25 | 0.25 | 0.25 | 0.25 |

***Supplementary Table 1: Nucleotide distributions used in superSTR simulation studies.*** The *P. falciparum GC content is sourced from the 3D7 isolate.*

| **Read Length** | Notes | *Homo sapiens 40.87% GC* | *Streptomyces coelicolor 72% GC* | *Plasmodium falciparum 13.5% GC* | *Uniform distribution 50% GC* |
|---|---|---|---|---|---|
| **16** | Not recommended, exercise caution | 1.31 | 1.25 | 1.19 | 1.31 |
| **36** | | 0.81 | 0.78 | 0.75 | 0.81 |
| **75** | Recommended read lengths for superSTR | 0.63 | 0.63 | 0.61 | 0.63 |
| **100** | | 0.56 | 0.55 | 0.54 | 0.56 |
| **125** | | 0.52 | 0.51 | 0.49 | 0.52 |
| **150** | | 0.49 | 0.48 | 0.47 | 0.49 |
| **200** | | 0.46 | 0.45 | 0.43 | 0.46 |
| **250** | | 0.44 | 0.42 | 0.40 | 0.44 |
| **300** | | 0.42 | 0.41 | 0.39 | 0.42 |

***Supplementary Table 2: Precomputed superSTR C thresholds for the four background distributions (Supplementary Table 1) used in simulation studies.***

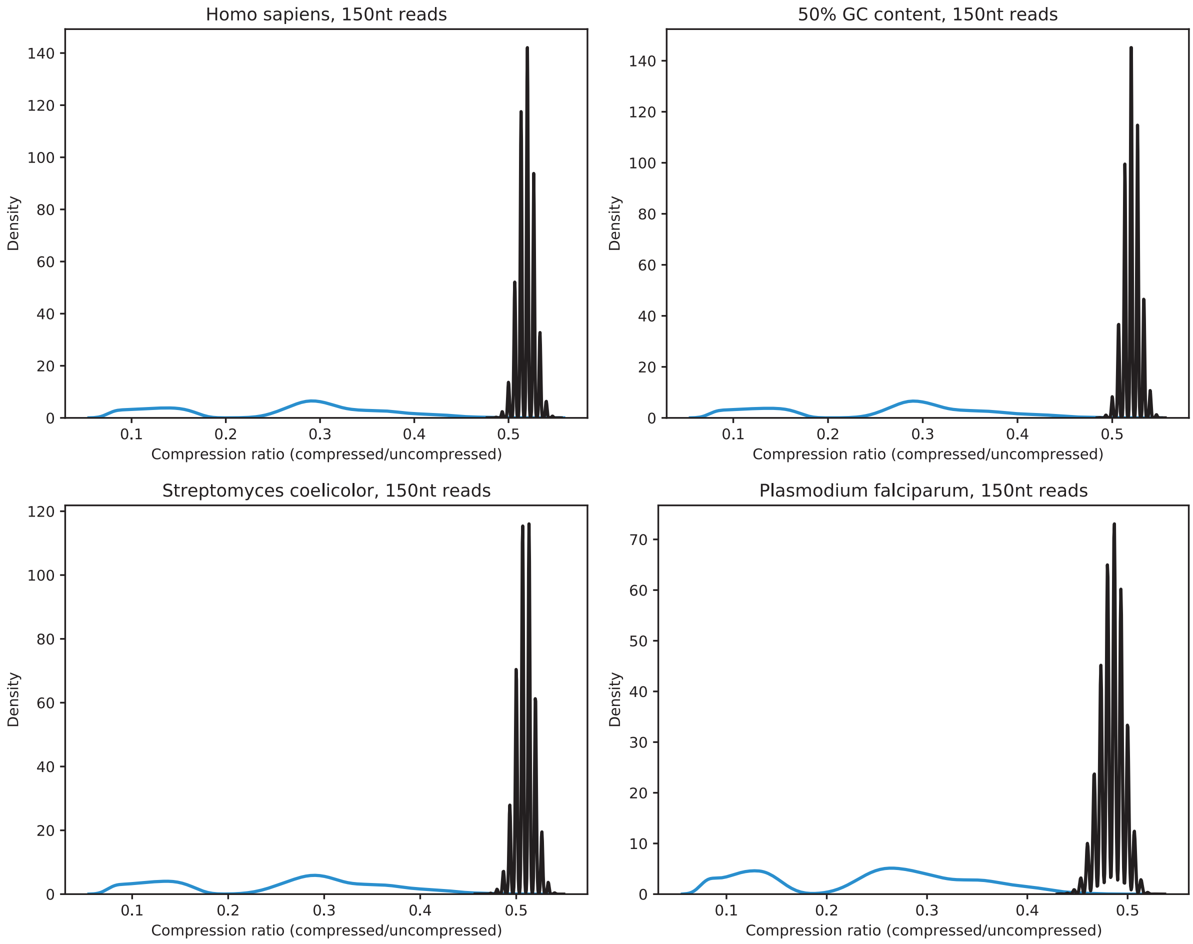


**Supplementary Figure S1: Distribution of compression heuristic values with fixed 150nt read length across four background nucleotide distributions**. We generated a total of 300,000 pseudorandom reads drawn from the indicated background distribution (shown in black). We then generated 300,000 repeat-containing reads; 100,000 pure repeat reads, 100,000 reads where the read contained a repetitive prefix or suffix, and 100,000 reads which contained a read with repeat sequence within the read flanked by pseudorandom sequence.

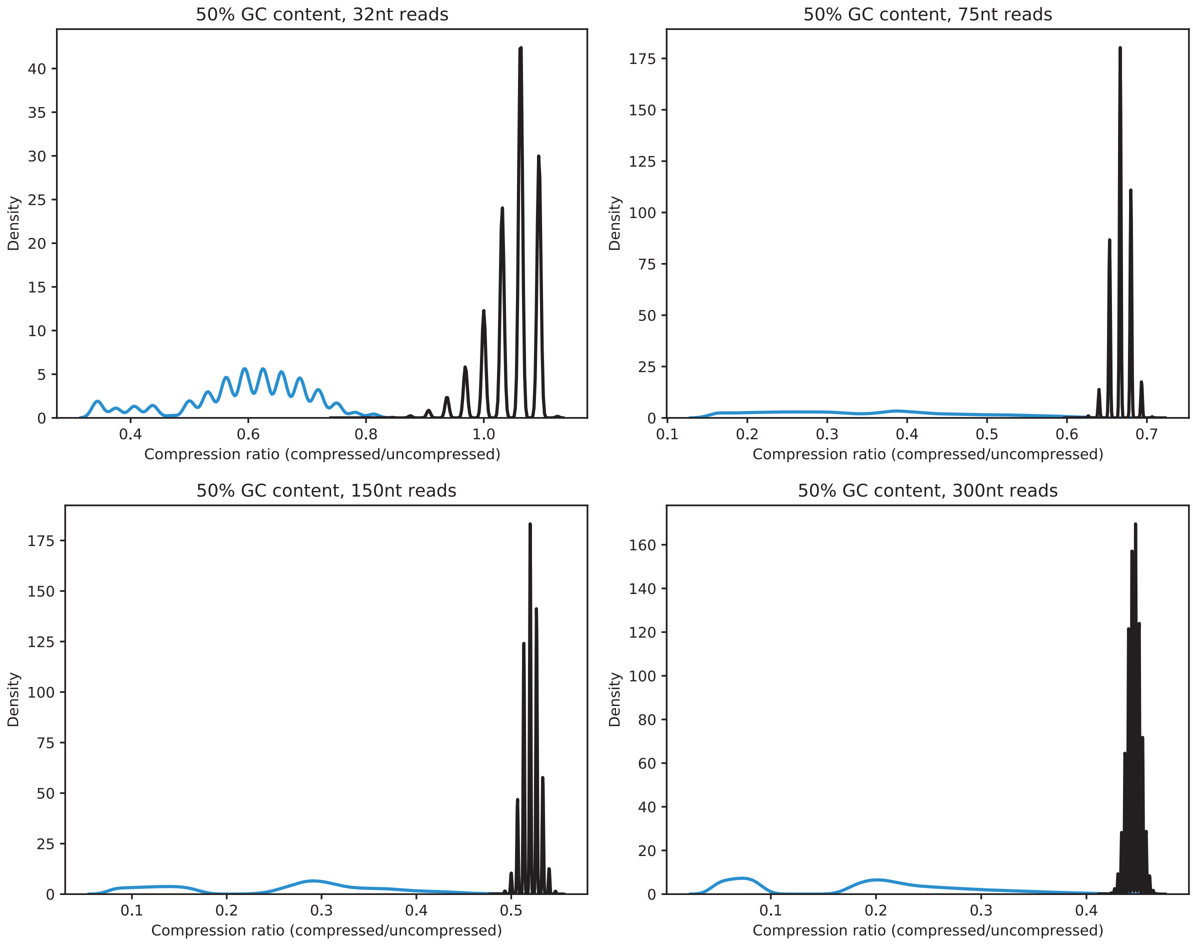


**Supplementary Figure S2: Distribution of compression heuristic values with varying read length drawn from a uniform background nucleotide distribution**. The uniform background nucleotide distribution is that with 50% GC content. Each plot shows 300,000 pseudorandom reads drawn from the background distribution (black) and 300,000 simulated repeat-containing reads. Note that in the 32nt example the pseudorandom draws exceed a compression ratio of 1.0 (as discussed in Supplementary Text).

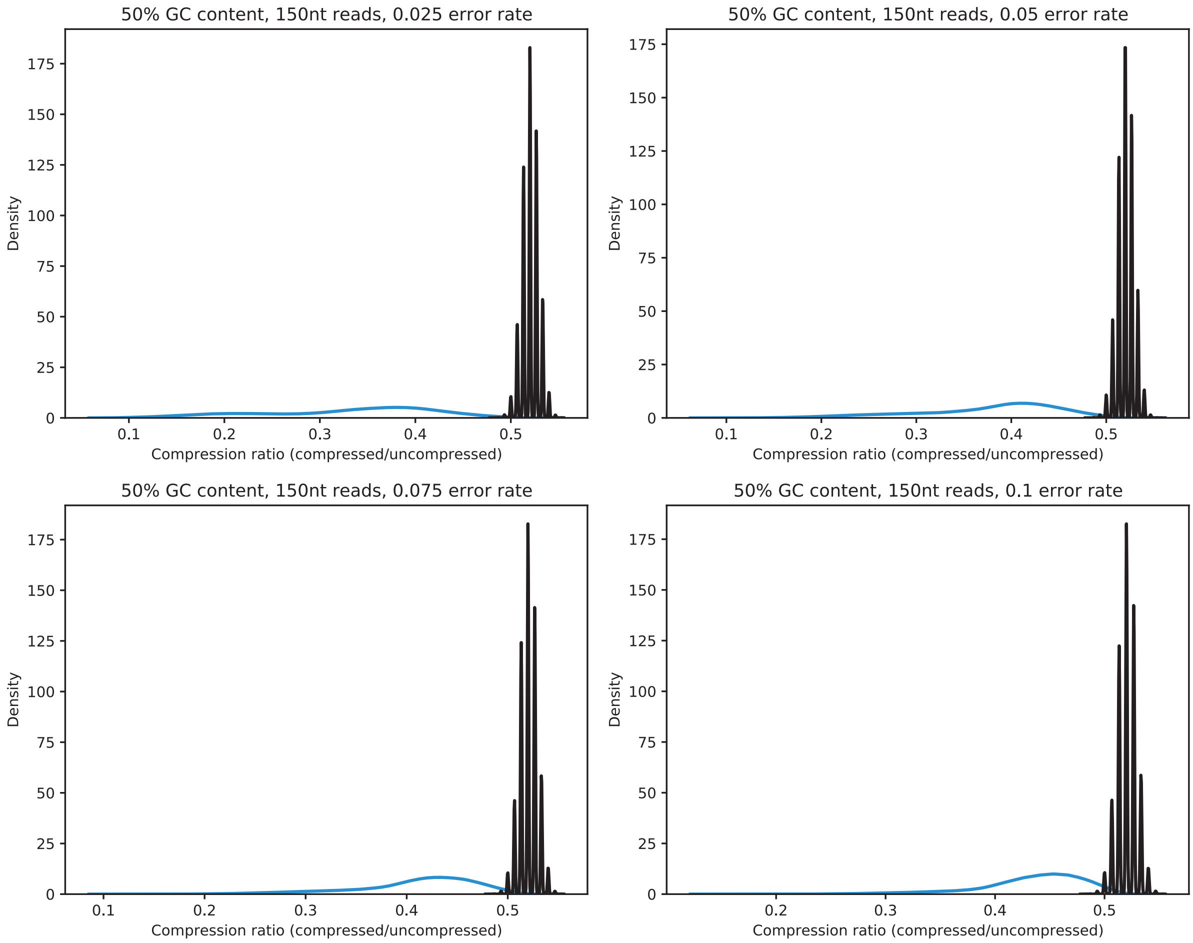


**Supplementary Figure S3: Distribution of compression heuristic values with 150nt read length drawn from a uniform background nucleotide distribution with error**. The uniform background nucleotide distribution is that with 50% GC content. Error rates were varied from 2.5% to 10% in 2.5% increments (note that the 0% error rate is shown in Supplementary Figure S1).

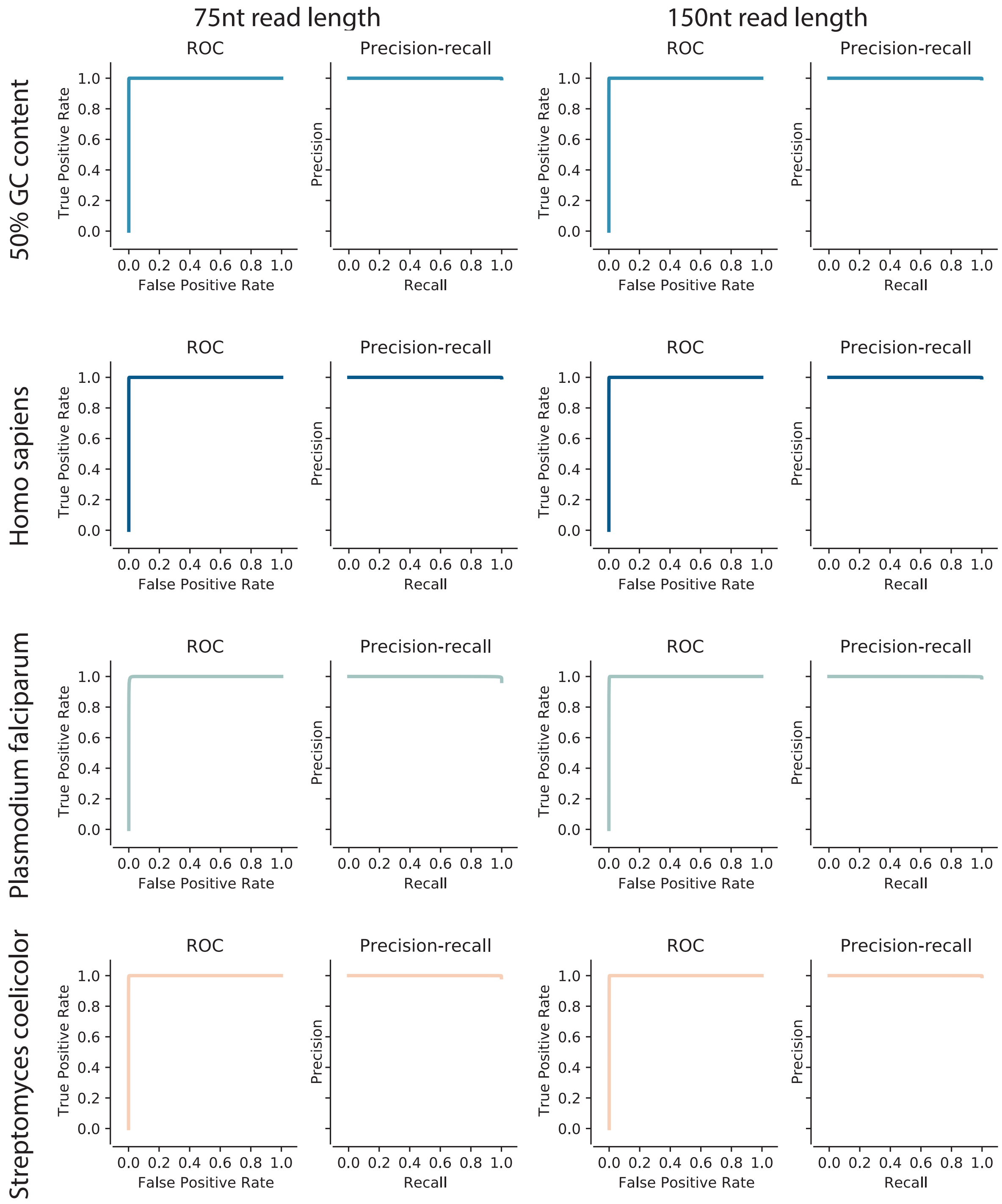


**Supplementary Figure S4: Receiver-operating characteristic and precision-recall curves for the compression heuristic in classifying 50,000 simulated reads as pseudorandom or repeat containing at 75nt and 150nt read lengths.** Area under ROC and PRC was greater than 0.99 in all displayed plots.

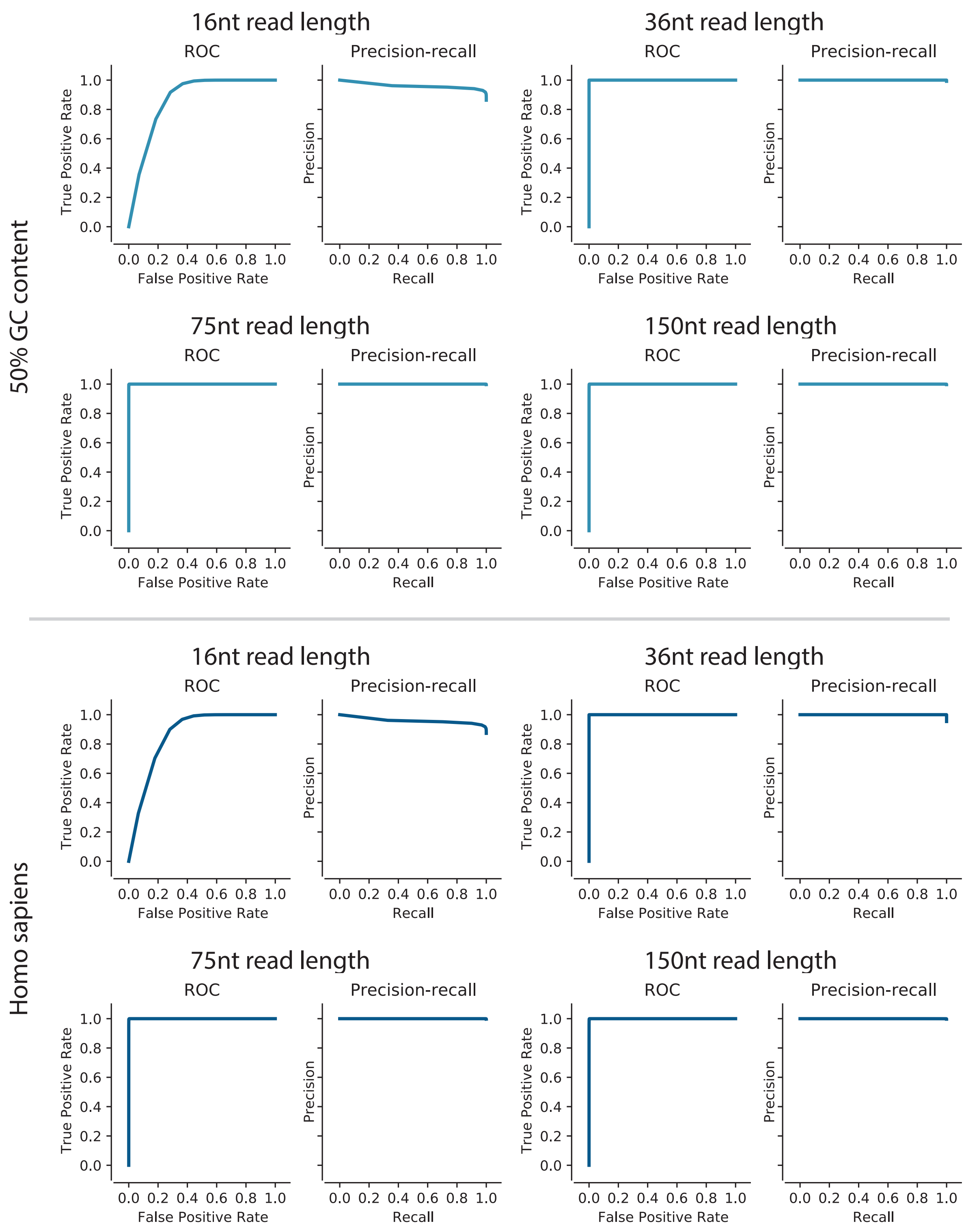


**Supplementary Figure S5: Receiver-operating characteristic and precision-recall curves for the compression heuristic in classifying 50,000 simulated reads as pseudorandom or repeat containing across extended read lengths for the Homo sapiens and uniform nucleotide distributions.**

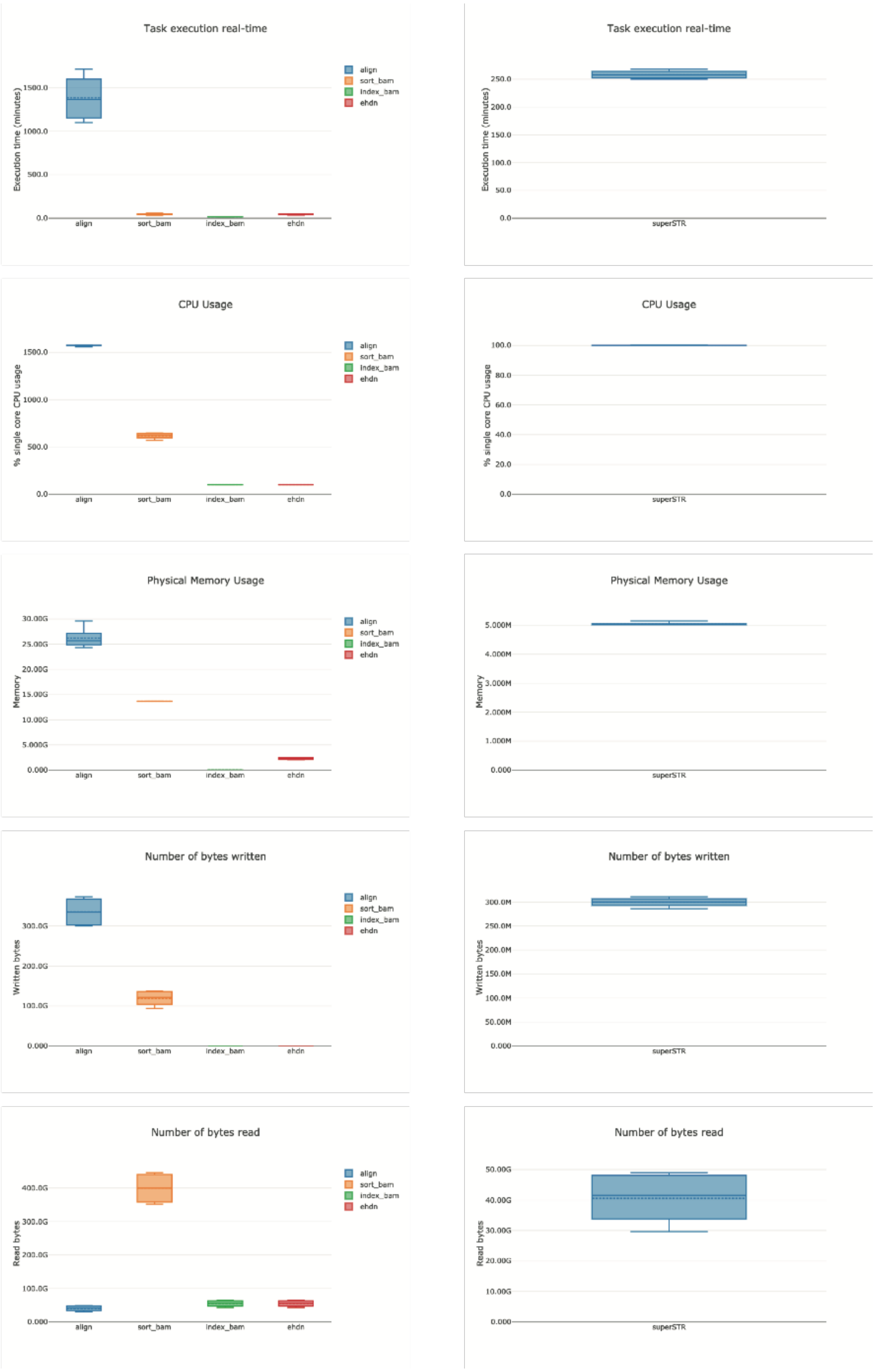


**Supplementary Figure S6: ExpansionHunter DeNovo/bwa-mem (left) and superSTR benchmarking (right) on WGS samples.**

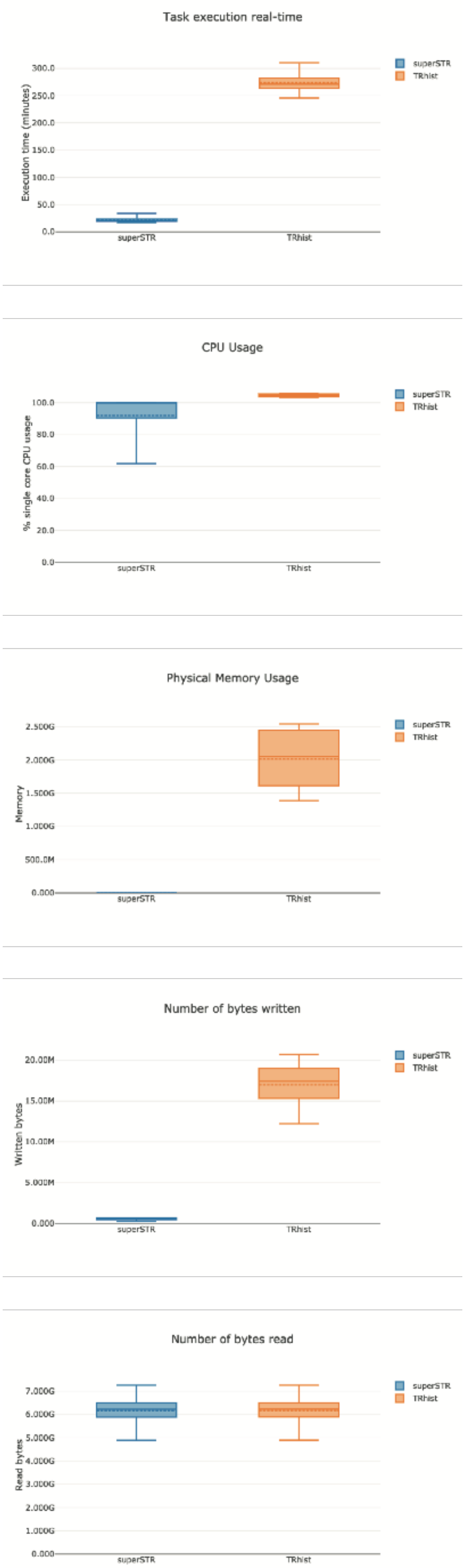


**Supplementary Figure S7: superSTR (left) and TRhist (right) benchmarking on RNA-seq samples.**
